# The non-transcranial TMS-evoked potential is an inherent source of ambiguity in TMS-EEG studies

**DOI:** 10.1101/337782

**Authors:** Virginia Conde, Leo Tomasevic, Irina Akopian, Konrad Stanek, Guilherme B. Saturnino, Axel Thielscher, Til Ole Bergmann, Hartwig Roman Siebner

**Author notes:** These authors contributed equally to this paper (Virginia Conde, Leo Tomasevic). These authors share senior authorship (Til Ole Bergmann, Hartwig Roman Siebner). Corresponding author: Hartwig R. Siebner, Head of Research, Danish Research Centre for Magnetic Resonance (DRCMR), Center for Functional and Diagnostic Imaging and Research, Copenhagen University Hospital Hvidovre, Kettegård Allé 30, 2650 Hvidovre, Denmark; http://www.drcmr.dk/siebner.

## Abstract

Transcranial Magnetic Stimulation (TMS) excites populations of neurons in the stimulated cortex, and the resulting activation may spread to connected brain regions. The distributed cortical response can be recorded with electroencephalography (EEG). Since TMS also stimulates peripheral sensory and motor axons and generates a loud “click” sound, the TMS-evoked EEG potential (TEP) not only reflects neural activity induced by transcranial neuronal excitation but also neural activity reflecting somatosensory and auditory processing. In 17 healthy young individuals, we systematically assessed the contribution of multisensory peripheral stimulation to TEPs using a TMS-compatible EEG system. Real TMS was delivered with a figure-of-eight coil over the left para-median posterior parietal cortex or superior frontal gyrus with the coil being oriented perpendicularly or in parallel to the target gyrus. We also recorded the EEG responses evoked by sham stimulation over the posterior parietal and superior frontal cortex, mimicking the auditory and somatosensory sensations evoked by real TMS. We applied state-of-the-art procedures to attenuate somatosensory and auditory confounds during real TMS, including the placement of a foam layer underneath the coil and auditory noise masking. Despite these precautions, the temporal and spatial features of the cortical potentials evoked by real TMS at the prefrontal and parietal site closely resembled the cortical potentials evoked by realistic sham TMS, both for early and late TEP components. Our findings stress the need to include a peripheral multisensory control stimulation in the study design to enable a dissociation between truly transcranial and non-transcranial components of TEPs.

## Introduction

Transcranial magnetic stimulation (TMS) produces a time-varying electric field that can directly excite neuronal populations in the cortical target area, bypassing the afferent sensory systems (Barker et al., 1985). The highly synchronized neural excitation of the target region spreads to inter-connected brain regions via the existing neuronal pathways which can then be captured with functional brain mapping techniques (Bergmann et al., 2016; Siebner et al., 2009). Electroencephalography (EEG) has been increasingly employed in recent years to measure the cortical responses evoked by focal TMS which, thanks to its excellent temporal resolution, can reveal how the local neural response spreads from the target site to functionally and structurally connected brain regions (Bergmann et al., 2016; Bortoletto et al., 2015; Ilmoniemi et al., 1997; Siebner et al., 2009).

TEPs vary in shape and number of components across cortical areas (Rosanova et al., 2009) and have been used to investigate cortical physiology both in health and certain neurological and psychiatric disorders (Farzan et al., 2016; Hallett et al., 2017; Kaskie and Ferrarelli, 2018; Massimini et al., 2012), with specific TEP components showing potential as clinical biomarkers (Manganotti et al., 2015). Connectivity measures have been derived from TMS-EEG data to infer how neuronal activations propagate across specific networks and how these networks change depending on different brain states (Bortoletto et al., 2015; Rosanova et al., 2009). The Perturbational Complexity Index (Casali et al., 2013), for example, reflects the spatiotemporal complexity of cortical responses to TMS and has been used as a connectivity marker for consciousness in humans (Rosanova et al., 2012). Moreover, TMS-EEG has been combined with pharmacological interventions to elucidate the mechanisms underlying the different TEP components (Darmani et al., 2016; Premoli et al., 2014a).

However, TMS does not only activate the human cortex transcranially. The time-varying electric field induces action potentials in myelinated axons in the extracranial tissue as well. Eddy currents evoked in the cerebrospinal fluid may also effectively stimulate proximal cranial nerves passing through foramina at the base of the skull (Schmid et al., 1995). Orthodromic action potential propagation in peripheral motor axons results in twitches of cranial muscles, which not only causes muscle potentials and electrode movement artifacts in the TEP recordings (Mutanen et al., 2013), but also a twitch-induced sensory input to the brain. When stimulating the motor cortex, re-afferent somatosensory stimulation also originates from the peripheral target muscles and contributes to TEP and TMS-induced oscillatory activity (Fecchio et al., 2017; Premoli et al., 2017).

In addition to causing peripheral somatosensory responses, the electrical discharge in the coil produces a loud “click” sound due to the mechanical quick expansion of the copper coil when the electric current passes through it, triggering auditory evoked potentials (Nikouline et al., 1999). Earplugs alone hardly attenuate even the airborne part of the “click”, but masking noise procedures can be used to minimize auditory co-stimulation. White noise or noise adjusted to the time-varying frequency of the TMS “click” can be administered via sound-proof in-ear headphones to prevent the TMS sound to be singled out by the brain (Massimini et al., 2005). Noise making can substantially reduce auditory evoked components in the TEPs, but often no complete suppression can be achieved at sound levels bearable for the participants, and a low frequency component can still be perceivable via bone conduction (Tchumatchenko and Reichenbach, 2014; ter Braack et al., 2015). A foam layer underneath the coil can dampen bone conduction and attenuate scalp sensations caused by mechanical coil vibration. However, the effectiveness of this method varies across participants (ter Braack et al., 2015).

At physiologically effective stimulus intensities, TMS will always cause significant peripheral costimulation, producing spatiotemporally complex cortical responses that do not result from direct transcranial cortical activation. The quantity and quality of somatosensory and auditory co-activation varies from site to site and depends on stimulation intensity and coil design. Since indirect multisensory (non-transcranial) and direct transcranial brain stimulation occur simultaneously, their evoked EEG responses are superimposed and hard to disentangle. Consequently, sham conditions have been used to characterize the non-transcranial multisensory contribution to the TEP. Sham stimulation is often achieved by the TMS coil being physically distanced from the scalp or tilted (Du et al., 2017; Fuggetta et al., 2008), thereby reducing the induced electric field in the cortex to a magnitude below stimulation threshold. The physical separation of the coil from the scalp preserves the airborne “click” sound but allows for little or no bone conduction and completely lacks somatosensory co-stimulation (Nikouline et al., 1999; ter Braack et al., 2015). The mere control by median nerve stimulation-evoked somatosensory potentials (Paus et al., 2001; Rosanova et al., 2009) not only lacks auditory stimulation but the evoked potentials may also not resemble those evoked by stimulating the scalp (Hashimoto, 1988). Sham TMS coils, generating only a very small electric field in the cortex, provide simultaneous somatosensory and auditory stimulation (Bonato et al., 2006; Opitz et al., 2014), but the area of stimulation is broader (Opitz et al., 2014) and somatosensory stimulation may be markedly reduced compared to real TMS (Bonato et al., 2006; Opitz et al., 2014). On the other hand, even when the transducing coil is placed on another body part such as the shoulder blade, stimulation still produced late evoked components reminiscent of those commonly seen in TEPs caused by real TMS (Herring et al., 2015).

This study systematically examines the contribution of multisensory co-stimulation to the TEP. We stimulated two different locations (frontal and parietal cortex) with two different coil orientations (orthogonal and parallel to the target sulcus/gyrus) and included a realistic sham condition for each location. The sham stimulation matched somatosensory and auditory co-stimulation of real TMS as closely as possible, while inducing only a subthreshold electric field in the brain. This enabled us to directly compare the EEG responses evoked by real and sham TMS. We hypothesized that non-transcranial multisensory co-stimulation makes a relevant contribution to TMS-evoked potentials. We therefore expected the spatiotemporal response patterns evoked by realistic sham and real TMS to resemble each other at both early and late latencies.

## Materials and methods

### Participants

The experiment was performed as part of a larger study investigating changes in connectivity during recovery from severe Traumatic Brain Injury (TBI) (see Conde et al. (2017) for more details). Seventeen healthy participants (10 females) with an age range from 19 to 31 years were included in the study of which 15 were completely naïve to TMS. The experimental procedures were approved by the Ethics committee of the Capital Region of Denmark (Region Hovedstaden). All participants gave informed written consent prior to the start of the experiment according to the declaration of Helsinki. Participants were asked to sit still and relax throughout the measurements while keeping their eyes open, and the chair was individually adjusted to achieve the most comfortable position by the use of arm, legs, and neck rests. None of the participants were using medication acting on the central nervous system by the time of the study. Information regarding hours of sleep, caffeine and tobacco intake, as well as levels of tiredness and discomfort (before and after the experiment, visual analogue scale from 0 to 10, 0 being lowest and 10 being highest level) was acquired via self-report.

### Experimental design

The experimental design is illustrated in Fig. 1. The experiment consisted of a single session per participant where both structural MRI and TMS combined with EEG (TMS-EEG) were performed. Structural MRI was always performed prior to TMS-EEG in order to acquire a T1-weighted image where the TMS target sites were individually identified and marked for online tracking by means of a frameless stereotactic neuronavigation system (Localite, St. Augustin, Germany).

**Fig. 1.**
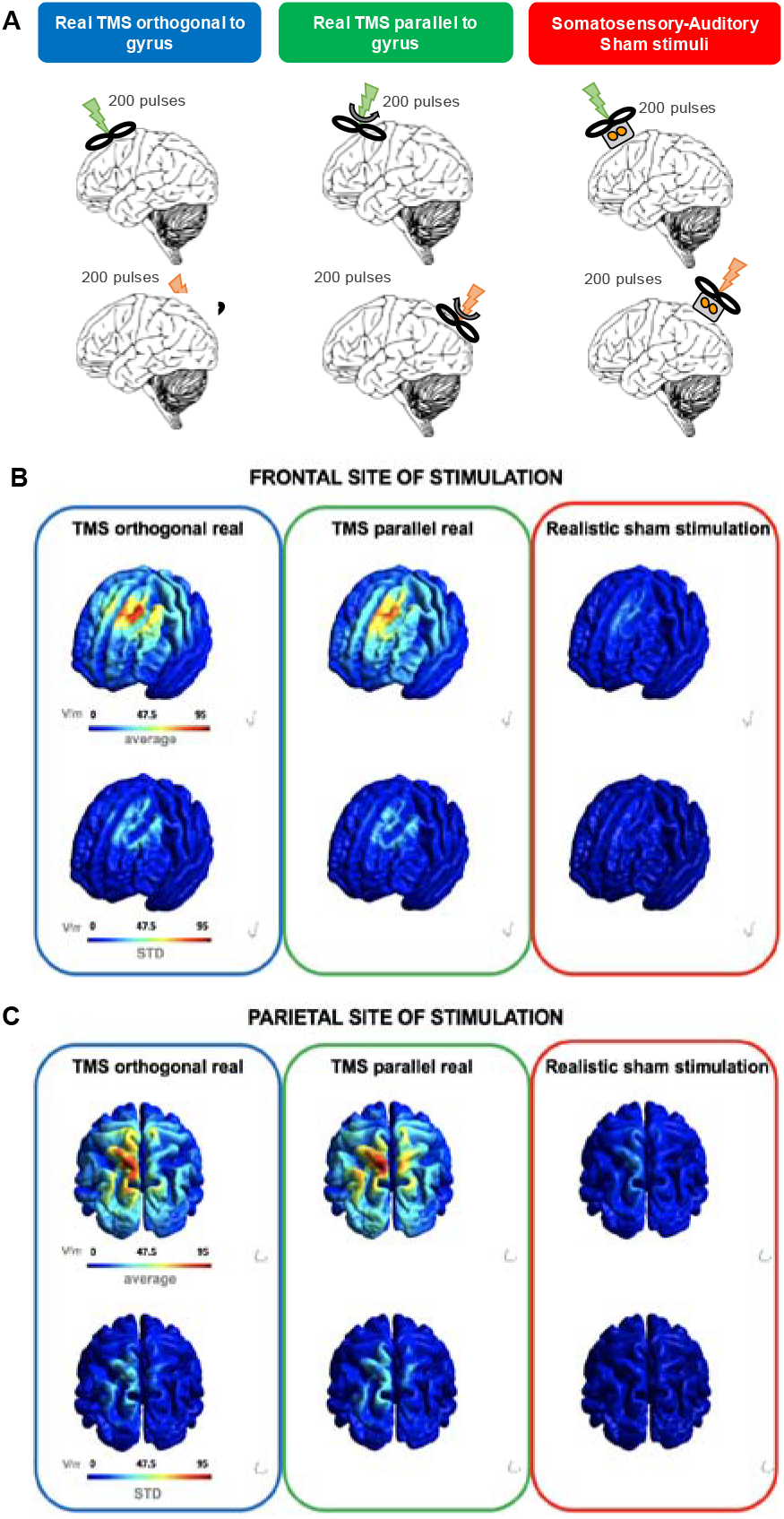
Experimental design and electric field modelling. **A**. Schematic representation of the experimental design showing the three different stimulation conditions and the two cortical target sites, namely left para-median superior frontal gyrus and left para-median superior parietal lobule. Curved arrows on the second column (TMS parallel) indicate the change of the coil angle with respect to the first column (TMS orthogonal) by 90 degrees (counter-clockwise for the frontal hotspot, clockwise for the parietal hotspot). A grey box with two orange circles inside (last column) represents the bipolar surface electrodes for electric stimulation. A total of 200 pulses were delivered per stimulation condition and cortical target. **B**. Group average of electric field maps for the frontal target site. **C**. Group average of electric field maps for the parietal target site. The color bar represents maximum strength of the electric field in V/m, ranging between 0 (blue), 47.5 (green-yellow), and a maximum of 95 (orange-red). Upper row shows the electric field strength maps across conditions. Bottom row shows the standard deviation of the strength maps per condition.

The experiment involved focal TMS of two brain areas (frontal and parietal cortex) with three stimulation conditions per cortical target site: two real TMS conditions with the coil oriented either perpendicular or in parallel to the cortical gyrus targeted by TMS and one somatosensory-auditory sham TMS (see Fig. 1).

The order of the six stimulation conditions was counterbalanced across participants, but always alternating between the prefrontal and parietal stimulation site in consecutive stimulation conditions. Two target sites were chosen for stimulation, namely the left para-median superior frontal gyrus (SPG) and left para-median superior parietal lobule (SPL) to enable conceptual within-study replication by targeting two para-median cortical areas. We chose these associative cortical areas because they are commonly targeted in TMS-EEG studies on disorders or consciousness (Marinazzo et al., 2014; Napolitani et al., 2014; Rosanova et al., 2009; Rosanova et al., 2012; Sarasso et al., 2014; Storm et al., 2017). In contrast to more lateral stimulation sites, TMS of these para-median areas elicits little to no muscle twitches. The determination of the exact coil position was based on individual anatomical MRIs as described previously (Rosanova et al., 2012).

For each stimulation site, we included two different coil orientations for real TMS. The longitudinal axis coil was either oriented orthogonally or in parallel to the orientation of the target gyrus referred to as “orthogonal” and “parallel” real TMS condition, respectively. The orthogonal coil orientation is considered to be the most optimal coil orientation to induce the strongest electric field within the target area (Thielscher et al., 2011). In the parallel real TMS condition, the coil orientation was rotated by 90 degrees relative to the orthogonal condition (counter-clockwise in the frontal hotspot and clockwise in the parietal hotspot due to physical constraints of the Neuronavigation system) with the longitudinal axis of the coil being aligned to the orientation of the gyrus.

We took state-of-the-art measures to reduce somatosensory and auditory TEP contamination. A thin layer of foam was placed under the coil and auditory noise masking was delivered throughout TMS measurements. Noise masking was delivered through in-ear headphones fitted inside foam earplugs (3M systems), and the masking noise was generated from the specific time-varying frequency of the coil as background noise with superimposed high-frequency coil “click” sounds as done previously (Herring et al., 2015). The sound pressure for noise masking was individually adjusted. The sound pressure was gradually increased (up to maximally 95 dB) until the participant could not hear the “click” sound of the coil with the TMS coil placed on their scalp or until they had reached their upper threshold for comfort.

After we had adjusted subjective stimulus intensity of electric somatosensory stimulation to the perceived intensity of orthogonal real TMS, each participant was asked to report the perceived focality of somatosensory stimulation on the scalp (defined as the extent of the area of scalp where the pulse was perceived, 10 being extremely narrow and 1 being extremely broad), the perceived loudness of the coil’s “click” sound, and the perceived overall discomfort related to stimulation after each of the six conditions. Participants were asked to give a score on a visual analogue scale (VAS) ranging from 0 to 10 with 0 corresponding to no perception and 10 to maximal perception.

### Magnetic Resonance Imaging

T1-weighted and T2-weighted structural images were acquired with a 3 Tesla TRIO Siemens scanner and a 16-channel head coil (Siemens Healthcare). Whole-brain T1-weighted and T2-weighted images were obtained with three-dimensional sequences (T1-w: TR = 2300ms, TE = 2.92ms, Voxel size = 1mm3 isotropic; T2-w: TR = 10000ms, TE = 52ms, Voxel size = 1×1×2 mm3 isotropic). T1-weighted images were acquired to individually identify and track the TMS hotspots with frameless stereotactic neuronavigation on each participant’s macrostructure. T2-weighted images were acquired together with T1-weighted images for the offline simulation of the induced electric fields in the cortex of each individual participant given the intensity of the stimulation and the distance of the coil from the scalp in each condition (SimNIBS 2.0, http://simnibs.de) (Thielscher et al., 2015)). Participants were provided with earplugs and instructed to lay still and to not move in between sequences.

### Electroencephalography

EEG was acquired with a TMS-compatible 64-channel system (Brain Products, 2 MR Plus 32-channel amplifiers) and a TMS-compatible EEG cap equipped with multitrodes (EasyCap, 61 equidistant multitrodes). Two multitrodes were used for EOG (below left eye, above right eye), reference was placed outside the cap on the forehead, and two ground multitrodes (one on the forehead, one over the left mastoid) were used to account for the long-lasting artifact induced by the cutaneous electric stimulation. Impedances of all electrodes were kept below 5 kOhm (Ilmoniemi and Kicic, 2010)Electrode leads were arranged orthogonal to the direction of the induced current in each condition in order to minimize TMS-induced artefacts (Sekiguchi et al., 2011).

Raw EEG signals were recorded with a sampling rate of 5 kHz at DC and only with an obligatory antialiasing low-pass filter of 1kHz (BrainVision Recorder, Brain Products, Munich, Germany). Baseline-corrected event-related potential (ERP) averages with common referencing were monitored online for visualization purposes. EEG electrode positions were digitized in each participant as an overlay of the 3D reconstructed scalp by means of a Neuronavigation system as reported previously (Herring et al., 2015). Participants were monitored to ensure that eyes were open, both by direct visual inspection and by identification of blinks in the raw EEG, and were touched at the hand in between TMS pulses as a signal to relax when muscle activity due to tonic contractions in cranial, jaw, or neck muscles was detected.

### Transcranial Magnetic Stimulation

Single-pulse TMS was performed with a PowerMAG 100 magnetic stimulator and a figure-of-eight shaped PMD45 coil with an outer winding diameter of 45 mm (Mag&More, Munich, Germany). The TMS pulse had a biphasic configuration and the first phase produced an inward current into the lateral wall of the targeted gyrus in the orthogonal condition. TMS intensity was individually adjusted for each cortical target site based on the local TEP response. Intensity of stimulation was kept the same between coil orientations within hotspots. We applied 50 TMS pulses at a jittered inter-trial interval (ITI) of 2 ± 0.4 s and measured the early (below 50 ms) peak EEG response using the average EEG signal from the channels neighboring the site of stimulation (Brain Products Visualizer, Munich, Germany). Starting with 60% of the maximum stimulator output (MSO), we gradually increased TMS intensity in steps of 2% MSO until TMS induced an early TEP peak with peak-to-peak amplitude of at least 6 μV. At this intensity, 200 pulses were delivered using the same jittered ITI as above.

### Somatosensory-auditory sham condition

The sham condition was designed to match the multisensory stimulation caused by real TMS as closely as possible (Rossi et al., 2007). Peripheral somatosensory stimulation was generated by cutaneous electric stimulation. We applied a square pulse delivered through bipolar electrodes (distance between electrodes: 25 mm), using a DS7A electric stimulator (Digitimer Ltd., Ft. Lauderdale, Florida, USA). The electrodes were attached to a plastic holder of 3.6 cm height, had a diameter of 8 mm and were previously soaked in water and fitted through holes cut in the fabric of the EEG cap. The electrical stimulus had a 200 μs duration and a maximum compliance voltage of 200 V. Please note that prior studies using bipolar electric stimulation with small electrode diameter report that a voltage of 330 and 2000 V is needed to stimulate the cortex (Cohen and Hallett, 1988; Merton and Morton, 1980). Intensity of the electrical stimulus was individually adjusted to match the sensation on the scalp induced by real TMS over each of the target hotspots via self-report of the participants.

The auditory stimulation was delivered through a TMS pulse with the figure-of-eight coil placed on top of the plastic holder of the bipolar electrode. The coil was placed directly on top of the plastic holder in order to retain optimal auditory stimulation levels (air and plastic/bone conduction) when compared to real TMS (Nikouline et al., 1999). The MSO of TMS was increased by 5% to account for the coil to scalp distance with regards to the strength of the auditory stimulation. The physical separation between coil and scalp ensured that no physiologically effective current was injected in the brain (Fig.1). Both, the cutaneous electrical and electromagnetic stimuli were delivered synchronously at ITIs that corresponded to the real TMS conditions using Signal software and a Micro 1401 system (Cambridge Electronic Design, Cambridge, UK).

### Data analyses

#### Visual analogue scales

Subjectively perceived focality of somatosensory stimulation, loudness of the “click” sound, and overall discomfort were recorded as individual scores on a VAS from 0 to 10. The self-reported scores were analyzed with R software (version 3.4.1.; https://www.r-project.org). Data distribution was explored by means of Q-Q plots as well as the Shapiro-Wilk normality test (data considered normally distributed if p > .05 and Q-Q plots show a linear fit). Since data were not normally distributed, Wilcoxon Signed-Ranked tests for dependent samples were used to contrast variables in a paired fashion, resulting in nine comparisons per stimulation site (three comparisons regarding focality, three regarding “click” sound, and three regarding discomfort, within and between hotspots per condition). Results were considered significant at p < 0.05 after Bonferroni-Holm correction for multiple comparisons (Ludbrook, 1998).

#### EEG pre-processing

The raw EEG data were pre-processed with in-house scripts programmed in Matlab (version 2016b, MathWorks, Natick, MA, USA). All the datasets were split in trial epochs starting 400 ms before and ending 1200 ms after the TMS pulse. The pulse artefact was removed from all datasets by interpolating the interval between −2 and 5 ms using cubic spline interpolation. Direct cutaneous electric stimulation over the scalp polarized the electrodes, which in turn resulted in a marked decay artefact affecting up to hundreds of milliseconds of signal. This decay artefact was removed by subtracting the best fit of a two-exponential function from each trial of each channel [42, 43]. Since TMS data were not affected by the decay artefact, this procedure was only applied to the sham datasets. Apart from decay artefact removal, all the analysis steps were identical for all datasets. The data were visually inspected and all the trials affected by strong artefacts (including eye-blinks) were discarded. If there were EEG channels with bad data quality, these channels were discarded and replaced by an interpolated signal using the weighted values of the surrounding channels. Finally, all datasets were filtered using a 50 Hz notch filter and a band-pass filter (high-pass: 1 Hz; low-pass: 80 Hz), down-sampled from 5 kHz to 500 Hz, baseline-corrected from −100 to −10 ms and re-referenced to the average of all electrodes. For each condition, the trials were averaged (constructing a grand-average) and the global mean field power (GMFP) was computed.

#### Analysis of the stimulation evoked EEG responses

We assessed the similarity of the evoked EEG data among the two real stimulation conditions (orthogonal and parallel coil orientation) and the realistic sham stimulation condition for each stimulation site (frontal and parietal) separately. We calculated the correlation between averaged temporal traces (correlation in time) and between potential distributions across channels (correlation in space) to evaluate the similarity between two stimulation conditions in time and in space, respectively. The temporal similarity was assessed channel by channel for the time interval ranging from 20 ms up to 410 ms after the TMS pulse. The very early post-stimulation time bin (< 20ms after stimulation) was not considered to avoid the first strong TMS and electric stimulation related artefacts [28, 36, 44]. Furthermore, we performed the same analysis on shorter intervals: early (parietal stimulation site: 20-58 ms; frontal stimulation site: 20-54 ms), middle (parietal: 58-144; frontal: 54-142 ms), and late response (parietal: 144-450 ms; frontal: 142-450). The three intervals were chosen based on the peaks observed in the GMFP of all subjects. The spatial similarity was evaluated for each time point by correlating the distribution of electrical potentials. In both cases, the correlation coefficients were estimated using the non-parametric Spearman method. To estimate the mean correlation among subjects both in time and space, the coefficients’ z-transform (Fisher’s z-transform) was averaged and subsequently inverse z-transformed. To assess the statistical significance of the correlation, we performed a pairwise t-test comparing the z-transformed coefficients before (−400 to - 10 ms) and after (20 to 410 ms) the stimulation. The significance level was set to < 0.05 after FDR correction for multiple comparisons.

#### Electric field simulations

Simulations of the electric fields generated by the TMS pulse for each participant and each condition were performed with SimNIBS software 2.0 (SimNIBS 2.0, http://simnibs.de) (Thielscher et al., 2015) using a realistic head model automatically generated from the individual T1-weighted and T2-weighted MR images as described elsewhere (Thielscher et al., 2015; Thielscher et al., 2011). In order to obtain average electric fields across subjects, the electric fields were interpolated in the middle of the segmented cortical gray matter, and transformed to the FSAverage template (Fischl, 2012), based on which the analysis was performed.

## Results

All participants underwent the measurements without reporting any adverse effects. On average, both tiredness and discomfort increased significantly from the beginning to the end of the experiment. The mean VAS scores for tiredness increased from 3.02 ± 1.78 to 6.20 ± 1.91 and for discomfort from 0.25 ± 0.71 to 1.85 ± 1.79, respectively. Participants reported an average amount of sleep of 7.25 hours during the night before the experiment (range from 6 to 9), ensuring that no participant was sleep-deprived by the time of the study. The intensity of magnetic stimulation in the real TMS conditions was 62.83 ± 4.94 % of MSO for the frontal hotspot, ranging from 53 to 75 % MSO, and 65.94 ± 3.62 % of MSO for the parietal hotspot, ranging from 60 to 72% MSO. For each target site, TMS was increased by 5% of MSO in the sham condition relative to the corresponding real TMS condition. The intensity of electric cutaneous stimulation was 9.25 ± 2.66 mA for the frontal hotspot (ranging from 2.5 to 13) and 10.88 ± 3.61 mA for the parietal hotspot (ranging from 5 to 21 mA).

While the intensity of the electric stimulation was individually adjusted to match the intensity of real TMS, the somatosensory perception was in all cases reported to be sharper (narrower area of the scalp) for sham than for real TMS conditions (Fig. 2). Accordingly, individual perception of focality differed significantly between all TMS conditions and their respective sham conditions (see below).

**Fig. 2.**
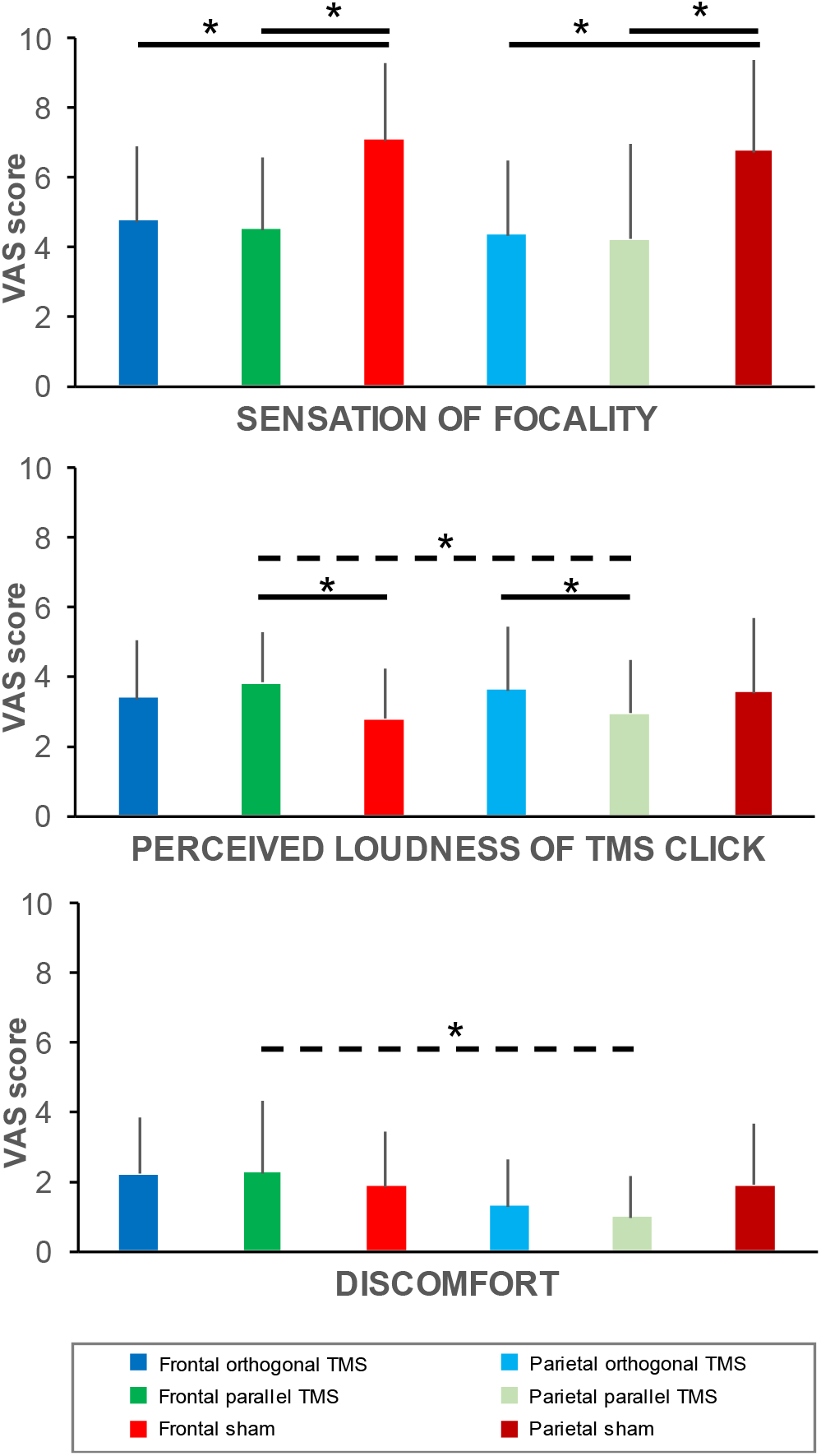
Self-reported perception of focality of stimulation (Upper panel), loudness of the perceived “click” sound (middle panel) and overall discomfort (lower panel). The columns represent the mean VAS scores (range: 0 to 10) and the error bars equal onefold standard deviation for each stimulation condition. The bold horizontal lines with an asterisk on top represent significant differences between two conditions for the same stimulation site (continuous lines) or between the frontal and parietal conditions (stippled line). Statistical comparisons used a Wilcoxon Signed-Ranked test with an alpha of 0.05/n (Bonferroni-Holm corrected for multiple comparisons).

Using a VAS with values ranging from 0 (no “click” perception) to 10 (maximal loudness of “click”), participants rated the intensity of auditory stimulation after each stimulation condition (Fig. 2). Only one participant in our study (one data point missing, n = 16 out of 17) reported complete absence of auditory perception of the “click” sound. All other participants reported VAS scores between 1 and 8, even though the volume of noise masking was adjusted to the noise level that most effectively attenuated the TMS-induced “click” sound in each participant without creating discomfort (ranging from 63 to 95 dB).

### Frontal stimulation site

For frontal stimulation, the perception of the “click” sound significantly differed between the parallel TMS condition and realistic sham condition (TMS parallel vs. Sham: p = 0.005; VAS = 3.82 ± 1.45 and VAS = 2.74 ± 1.49 respectively), but not for the commonly used orthogonal TMS condition (TMS orthogonal vs. Sham: p = 0.28). There were no significant differences between the two real TMS conditions in perception (TMS orthogonal vs. TMS parallel: focality p = 0.98; “click” sound p = 0.15; discomfort p = 0.76). Focality was significantly different between each real TMS condition and sham (TMS orthogonal vs. Sham: p = 0.003; TMS parallel vs. Sham: p = 0.002; TMS orthogonal VAS: 4.71 ± 2.17; TMS parallel VAS: 4.5 ± 2.06; Sham VAS = 7.09 ± 2.24). Finally, discomfort was not significantly different between real TMS conditions and sham (TMS orthogonal vs. Sham: p = 0.23; TMS parallel vs. Sham: p = 0.36).

### Parietal stimulation site

For parietal stimulation, a difference in “click” perception between conditions was only present when comparing both real TMS conditions (orthogonal and parallel) (TMS orthogonal vs. TMS parallel “click” sound p = 0.01; TMS orthogonal VAS: 3.59 ± 1.85; TMS parallel VAS: 2.91 ± 1.59), but not when the realistic sham condition was compared with the two real conditions (TMS orthogonal vs. Sham “click” sound p = 0.42; TMS parallel vs. Sham “click” sound p = 0.29). Focality was significantly different between real TMS conditions and sham only (TMS orthogonal vs. Sham: p = 0.005; TMS parallel vs. Sham p = 0.004; TMS orthogonal vs. TMS parallel: p = 0.93; TMS orthogonal VAS: 4.34 ± 2.12; TMS parallel VAS: 4.21 ± 2.73; Sham VAS: 6.74 ± 2.60). Perceived discomfort was not significantly different across any condition (TMS orthogonal vs. TMS parallel: p = 0.15; TMS orthogonal vs. Sham discomfort p = 0.09; TMS parallel vs. Sham: p = 0.06).

### Frontal versus parietal stimulation

Focality, “click” sound, and discomfort were not perceived to be different across hotspots when comparing the orthogonal TMS conditions between the frontal and parietal hotspots (focality: p = 0.49; “click” sound perception: p = 0.34; discomfort: p = 0.034). This was different for the parallel real TMS condition, in which “click” sound perception (p = 0.01; TMS parallel frontal VAS: 3.82 ± 1.46; TMS parallel parietal VAS: 2.91 ± 1.59) and discomfort (p = 0.005; TMS parallel frontal VAS: 2.64 ± 2.03; TMS parallel parietal VAS: 0.94 ± 1.18) were reported to be significantly different between the frontal and the parietal stimulation sites, in contrast to the perceived focality (p = 0.34). Subjective ratings for the sham stimulation were not significantly different between the frontal and parietal hotspots (focality: p = 0. 20; “click” sound: p = 0.17; discomfort p = 0.86).

#### Stimulation-evoked EEG responses

Both the grand-average of TEPs and GMFP showed significant similarity between real and sham stimulation in the temporal domain. The realistic sham condition evoked a response profile in the EEG that shared the timing and spatial distribution of major EEG peaks evoked with real TMS. Providing internal replication, this similarity was observed for the frontal and parietal stimulation site (Fig. 3 and 4).

**Fig. 3.**
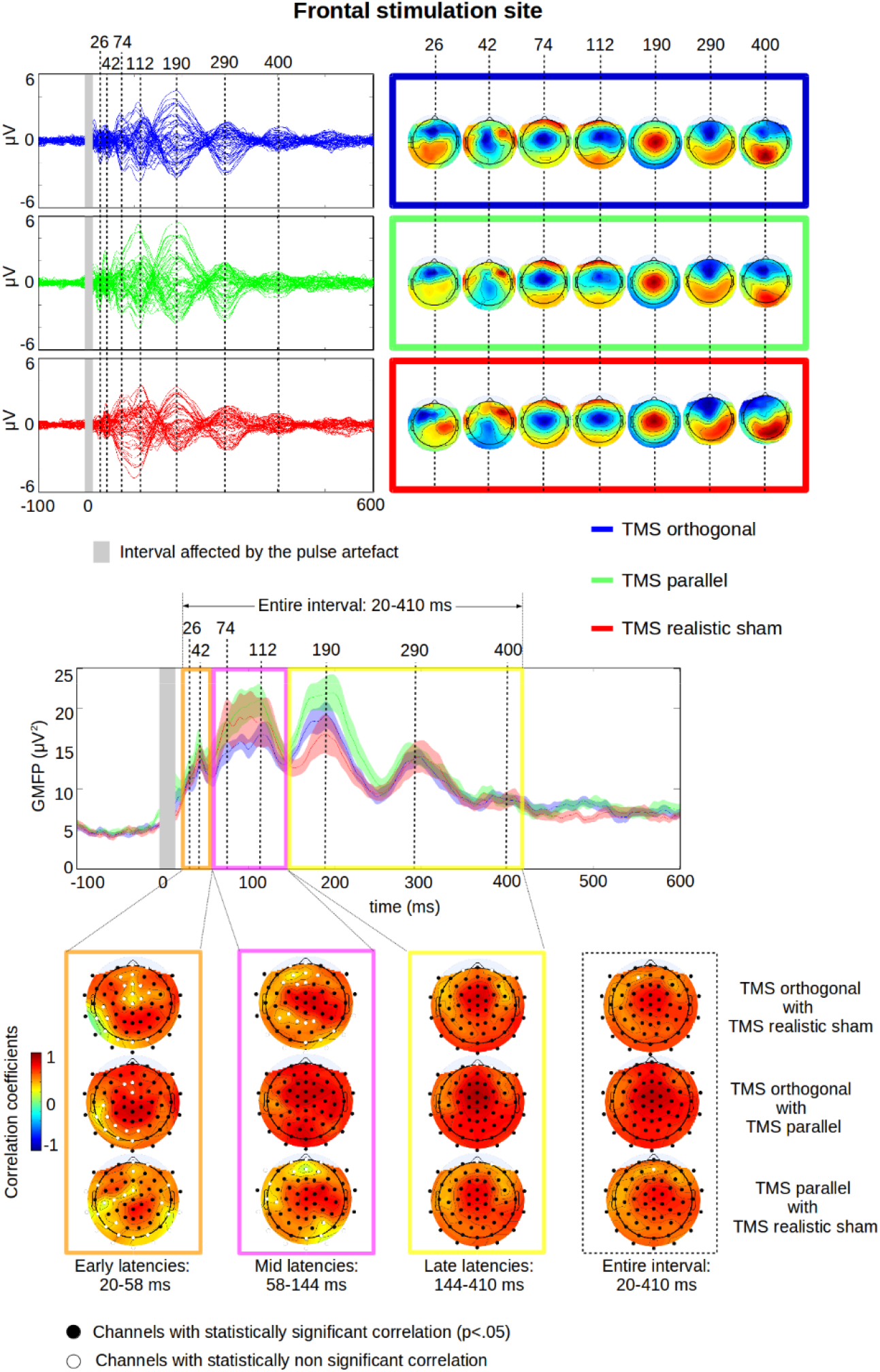
Group EEG data evoked by stimulation targeting the left paramedian frontal cortex. Upper panel: Grand averages of TEPs for each EEG channel and the topographic distribution of the electric potentials of the identified peaks (maps on the right). The responses evoked by TMS delivered orthogonal to the target gyrus are presented in blue colour. The responses evoked by TMS delivered with the coil oriented in parallel to the target gyrus are presented in green colour. The responses evoked by the somatosensory-auditory sham stimulation are labelled in red. Middle panel: Global mean field power (GMFP) of the three stimulation conditions and the selection of intervals for the time correlation (early, middle, late response and the entire interval. Lower panel: Topographic distribution of the average correlation in the selected intervals between two of the three conditions (orthogonal real TMS vs realistic sham TMS; orthogonal real TMS vs. parallel real TMS; parallel real TMS vs. realistic sham TMS). Only a few channels marked as white dots showed no statistically significant correlation between conditions.

**Fig. 4.**
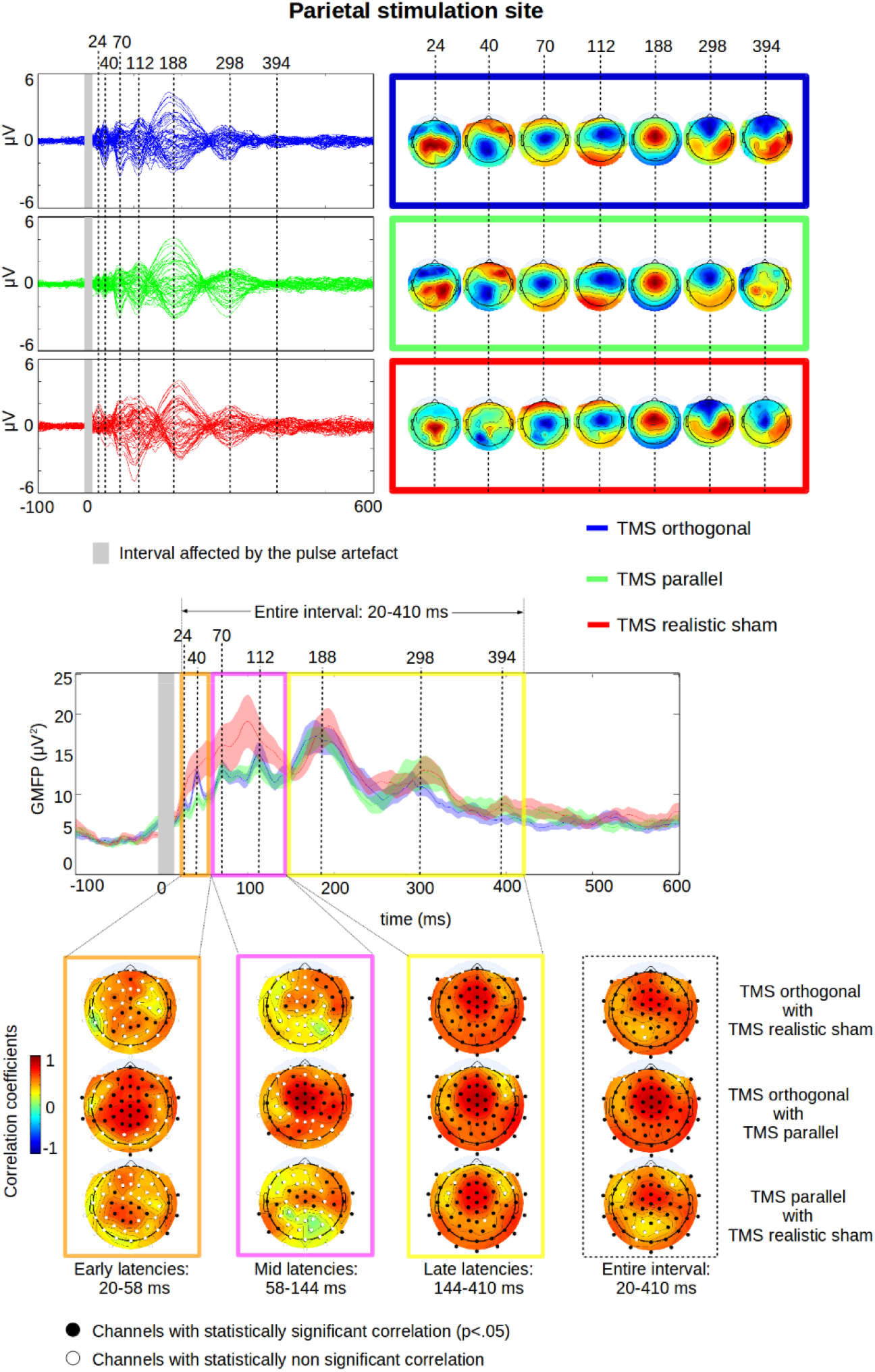
Group EEG data evoked by stimulation targeting the left paramedian parietal cortex. Upper panel: Grand averages of TEPs for each EEG channel and the topographic distribution of the electric potentials of the identified peaks (maps on the right). The responses evoked by TMS delivered orthogonal to the target gyrus are presented in blue colour. The responses evoked by TMS delivered with the coil oriented in parallel to the target gyrus are presented in green colour. The responses evoked by the somatosensory-auditory sham stimulation are labelled in red. Middle panel: Global mean field power (GMFP) of the three stimulation conditions and the selection of intervals for the time correlation (early, middle, late response and the entire interval. Lower panel: Topographic distribution of the average correlation in the selected intervals between two of the three conditions (orthogonal real TMS vs realistic sham TMS; orthogonal real TMS vs. parallel real TMS; parallel real TMS vs. realistic sham TMS).

The similarity of the EEG response to real and sham TMS was confirmed by a significant temporal correlation of the evoked potentials over the entire 20-450 ms interval expressed in the majority of EEG channels (Fig. 3 and 4). When the post-stimulation period was split into early, middle, and late poststimulation intervals, the widespread correlation of the temporal EEG response at a given channel was found for all three intervals, including the relevant peak responses (Fig. 3 and 4). The strength of the temporal correlation was spatially less pronounced at shorter post-stimulus intervals. The electrodes with the highest degree of correlation clustered in the central region, corresponding to the location that maximally represented the electric potentials identified at the peaks of the GMFP (Fig. 3 and 4).

We also found a strong similarity of the stimulation-evoked responses in the spatial domain, with real and sham conditions being closely matched in terms of the evoked spatial response pattern. The similarity between the spatial distributions of evoked responses over the scalp was confirmed by a correlation analysis that compared the site-specific real and sham conditions using the mean EEG amplitude of each individual at each post-stimulation time point across all electrode sites (Fig. 5). For the frontal target site, the spatial correlations of the EEG response between sham and real TMS conditions were significant over the entire post-stimulation period from 20 ms to 410 ms after stimulation (Fig. 5, upper panel). For the parietal stimulation site, the spatial correlations of the EEG response between sham and real TMS were significant for most of the tested interval, interrupted by short periods where correlation did not reach significance (Fig. 5, lower panel). The peaks at which spatial correlations between real and sham stimulation reached relative maxima corresponded to the timing of the peaks identified by GMFP, showing that the majority of the power was expressed by similar electrodes and at similar time points for real and sham TEPs (Fig. 5).

**Fig. 5.**
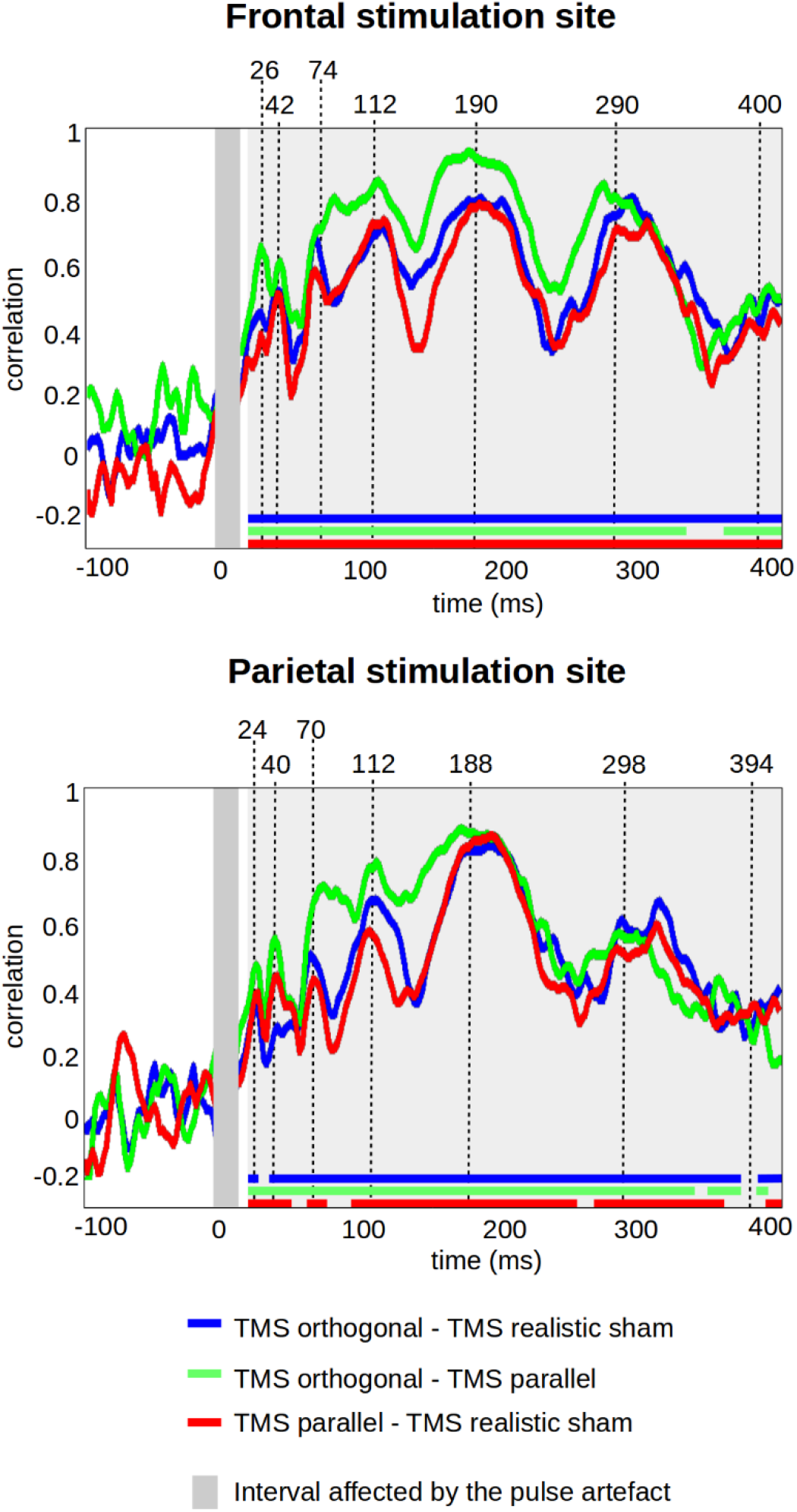
Spatial similarity of cortical responses evoked by real and sham TMS. The correlation between the distribution of the potentials over the scalp in different conditions (orthogonal TMS/sham; orthogonal TMS/tangential TMS; tangential TMS/sham) for the frontal (upper) and parietal (lower) stimulation spot. At the bottom of each figure the statistically significant time intervals are shown for each correlation analysis as a bold timeline. The interruptions indicate periods during which correlation did not reach significance. With a few exceptions, spatial correlations were significant between conditions across the entire post-stimulation interval.

We simulated the induced electric fields in each subject to estimate the residual electromagnetic stimulation in the sham condition and to compare the estimated values to those induced by the two real TMS conditions. The average and standard deviation of the electric field distribution is illustrated in Fig. 1. The maximum electric field strength averaged over all participants was comparable across real TMS conditions and well above the reported threshold for neuronal activation as recorded by EEG (> 50 V/m), indicating effective cortical stimulation in the real TMS conditions (Casali et al., 2010; Casali et al., 2013; Massimini et al., 2005; Rosanova et al., 2009; Sarasso et al., 2015). For the frontal site, mean peak electric field strength was 94 V/m for the real TMS condition in which TMS induced a field that was oriented orthogonally to the target gyrus, and 86.7 V/m for a parallel coil orientation. At the parietal site, mean peak electric filed strengths were 78.8 V/m for orthogonal TMS and 83.4 V/m for parallel TMS. The peak electric field strength was reduced by a factor of 4.24 (frontal) and 3.51 (parietal) with respect to real TMS (Frontal sham: 22.81 V/m; Parietal sham: 22.45 V/m), inducing a peak electric field strength well below the threshold to excite cortical neurons.

## Discussion

In the present study, we report findings that indicate that non-transcranial multisensory co-stimulation makes a substantial contribution to TEP components commonly interpreted as the direct brain’s response to the electric field induced by transcranial magnetic stimulation. When conducting combined TMS-EEG recordings, even state-of-the-art auditory noise masking and foam padding achieve only imperfect suppression of both the TMS “click”-related auditory input and the somatosensory input evoked by inductive electric stimulation of myelinated peripheral nerve axons. This is the first study to our knowledge that systematically assessed the impact of this multisensory co-stimulation on the EEG activity evoked by focal TMS targeting non-motor prefrontal and posterior parietal cortex. Although we implemented state-of-the art measures to attenuate multisensory co-stimulation, the cortical potentials evoked by real and sham TMS at the prefrontal and parietal site closely resembled each other, both in temporal shape and spatial distribution. This similarity might be even greater than the one shown in the present study, because our realistic sham condition did not perfectly match the multisensory input evoked by TMS in the somatosensory domain. The close resemblance of EEG responses evoked by real TMS and realistic sham stimulation shows that the non-transcranial TEP is an inherent source of ambiguity in TMS-EEG studies. Therefore, future TMS-EEG studies need to actively show that multisensory costimulation was suppressed completely. This could be achieved by showing that participants perform at chance level in a two-alternative forced choice test in which they indicate whether they have received TMS or not. If participants still can dissociate between TMS and no-TMS trials after all measures are taken to suppress multisensory co-stimulation, the experimental design needs to include a realistic sham control condition which mimics multi-sensory co-stimulation as closely as possible.

### Peripherally evoked potentials evoked by multisensory stimulation

Although our realistic sham stimulation did not perfectly match the multisensory input associated with real TMS, the temporal and spatial patterns of the peripherally-evoked cortical responses closely resembled the spatiotemporal patterns of TEPs evoked in the real TMS conditions. In the temporal domain, evoked peak latencies closely matched the TEP latencies evoked by real TMS at early, middle, and late post-stimulation intervals. Peak correspondence was found 40-400 ms post stimulation for the frontal target site and 70-400 ms for the parietal target side, including the classic N100 central negativity often reported in TMS-EEG studies (Du et al., 2017). Likewise, the topographical distribution of the evoked responses showed a significant correlation between sham and real TMS conditions for almost the entire 20-410 ms post-stimulation time window. Using a sham condition that consisted of real TMS delivered to the shoulder, Herring et al. (Herring et al., 2015) showed that sham stimulation induced a cortical response pattern that was similar to the one evoked by real TMS over the scalp, primarily at late peak latencies (> 80 ms post stimulation). Extending these findings, we show that concurrent cranial somatosensory and auditory stimulation mimicking TMS contributes substantially to the TEP also at early latencies.

The similarity between realistic sham and real TMS between 20 and 80 ms after TMS can be attributed to the auditory and somatosensory features of the realistic sham condition. Firstly, inductive electric stimulation of somatosensory nerve fibers in the skin underlying the TMS coil resulted in early cortical responses which could be due to somatosensory evoked potentials (SEP). Indeed, early SEP components are already present 15 ms after peripheral trigeminal stimulation (Malcharek et al., 2011). Peripheral trigeminal stimulation has also been shown to modulate the amplitude of motor evoked potentials triggered by TMS of the motor hand area, starting 40-50 ms after peripheral stimulation (Siebner et al., 1999). Of note, the parasagittal dura mater contains myelinated fast-conducting A-beta fibers. These fibers have most likely been stimulated by real TMS in a coil orientation-dependent fashion, contributing to trigeminal somatosensory stimulation in the real TMS conditions. Inductive electric stimulation of motor nerve fibers, especially peripheral branches of the facial nerve, might have caused secondary sensory input through the induction of muscle twitches, especially for the frontal target site. Secondly, residual “click” sound perception of the TMS pulse might have evoked mid-latency peaks of the auditory evoked potentials (AEP) which are expressed already 20 ms after stimulation on the scalp (Holt and Ozdamar, 2016). A recent study showed that AEPs are reliably evoked by very short gaps during noise stimulation (Alhussaini et al., 2018). Hence, the transient “click”-induced modulation of acoustic input might effectively evoke AEPs, even in the context of a noise stimulation background.

We also found a close resemblance of the EEG response between sham and real TMS stimulation conditions for the later components evoked by both realistic sham and real TMS, including the N100 and P180 components, commonly described as the N1-P2 complex for both auditory and somatosensory stimulation (Goff et al., 1977; Hyde, 1997). The auditory N1-P2 peaks at frontocentral scalp electrodes as a result of respectively oriented dipoles in bilateral temporal cortices (Zouridakis et al., 1998), and somatosensory components at > 100 ms originate from bilateral secondary somatosensory cortices (Allison et al., 1992). The N100 is of particular interest as has been associated with GABA-B-ergic inhibition based on pharmacological interventions (Premoli et al., 2014a) and paired-pulse TMS (Opie et al., 2017; Premoli et al., 2014b; Rogasch et al., 2012), as well as by its amplitude correlation with the silent period duration (Farzan et al., 2013). Notably, Du et al. (Du et al., 2017) observed a vertex N100 of similar amplitude after TMS of prefrontal, motor, primary auditory cortices, vertex, and cerebellum, and concluded that the N100 is a ubiquitous TEP reflecting a general property of the cerebral cortex. Our findings point rather to the conclusion that the N100 observed over the vertex is at least to a great extent a non-transcranial sensory evoked potential.

### Implications for studies of transcranial evoked potentials

The close resemblance of TMS and sham-evoked potentials does by no means imply that specific TEP components can be always and fully explained by multisensory-evoked potentials. On the contrary, TEP recordings hold great potential for probing the local and distributed brain response to focal TMS. Since the multisensory components overlap substantially with the truly transcranial components, it is necessary to disentangle the multisensory temporal and spatial response patterns from the truly transcranially-evoked brain response. The true TEP components may become only evident after subtraction of the multisensory components or in experimental designs that effectively account for multisensory stimulation as a confound. In the study of Herring et al. (Herring et al., 2015), for instance, the authors found a left occipital N40 component following left visual cortex TMS but not multisensory sham that can hardly be explained by somatosensory or auditory co-stimulation. If the topography of a TEP component is clearly lateralized and confined to the stimulation site, such component is less likely to be the mere result of multisensory stimulation which often shows a different voltage distribution. Also, the GABA-B-receptor-mediated amplitude modulation of an N100 component lateralized to the stimulated left sensorimotor cortex most likely reflects a local cortical effect at the target site (Premoli et al., 2014a). In contrast, GABA-A receptor-mediated amplitude modulations of the TEP have been reported to only be significant in the hemisphere contralateral to stimulation (Premoli et al., 2014a), and future work has to clarify the degree to which remote effects like this are due to distant scalp projections of a local dipole, a network spread of transcranially-induced activity, or pharmacological effects on multisensory cortical processing. Studies using similar GABA-mediating drugs such as benzodiazepines have consistently reported effects on AEPs and SEPs also at 100 ms, reinforcing the need to further investigate the purely transcranial effects of drugs on the TEP (Abduljawad et al., 2001; Scaife et al., 2006). Our findings are compatible with the notion that local activations at the target site may predominantly arise from transcranial stimulation particularly in the early post-stimulation period. For electrodes close to the stimulated region, the similarity between sham and real TMS was less consistent. The stronger dissimilarity of evoked responses 24-70 ms after stimulation may thus be due to the local activations after real TMS as compared to sham. Alternatively, this dissimilarity may have resulted from methodological issues since the decay artefacts resulting from transcutaneous electric stimulation were also strongest at the stimulation site, and the early post-stimulation interval included less time points than the middle or late post-stimulation intervals potentially decreasing similarity between stimulation conditions.

In a recent study aiming to disentangle the cortical origin of TEPs, Gosseries et al. targeted both lesioned and preserved cortical tissue in two patients with unresponsive wakefulness syndrome and multi-focal brain injury (Gosseries et al., 2015). In these patients, TEPs were completely absent when TMS directly targeted the lesioned cortex, whereas TEPs were preserved when targeting non-lesioned cortex, keeping multisensory co-stimulation comparable (Gosseries et al., 2015). These results show that a local cortical response can be evoked by TMS, but does not rule out a substantial multisensory contribution to TEPs recorded in healthy conscious individuals. It should also be noted that the first patient had additional brain stem lesions in the pons, medulla and cerebellar peduncles. These additional lesions might have blocked peripheral somatosensory input from the lesioned but not from the non-lesioned hemisphere. The second patient had massive bilateral hemispheric lesions, involving auditory and somatosensory cortex bilaterally. Again, this might have prevented the occurrence of cortical responses caused by multisensory co-stimulation. It also seems that substantially higher TMS intensities were applied by Gosseries et al. and the local responses had much larger amplitudes than those normally obtained in healthy conscious individuals. Finally, in patients with disorders of consciousness it is not possible to individually adjust the sound pressure of the noise masking, potentially resulting in higher sound pressures than those tolerated by healthy individuals.

### Can auditory and somatosensory stimulation be completely suppressed in awake individuals without brain lesions?

The evidence obtained in unresponsive patients with massive multi-focal brain damage (Gosseries et al., 2015) cannot be generalized to other studies and does not imply that those components are principally of transcranial origin when observed under different conditions. Special care needs to be taken when contrasting different physiological states (e.g., drug challenges, vigilance or attentional states, etc.) or groups (e.g., psychiatric or neurological patients) for which also a modulatory effect on auditory or somatosensory evoked potentials is conceivable or in some cases known. It has been proposed that multisensory co-stimulation does not account for any TEP components as long as both auditory and somatosensory perception are suppressed by noise masking and foam padding (Gosseries et al., 2015). Unfortunately, a complete suppression is often not achievable when studying fully awake individuals, even when following best practice procedures as reported in the present study. We implemented all measures currently advised to attenuate multisensory co-stimulation (i.e., individualized noise masking, foam padding, and stimulation sites close to the midline) and still observed multisensory evoked potentials, while almost all participants reported residual auditory and tactile perception of the TMS pulses. Unlike in other studies for which complete suppression of TMS “click” sound perception has been reported (Casula et al., 2017; Gosseries et al., 2015; Massimini et al., 2005), we systematically asked participants to rate perceptual intensity after each stimulation condition. Only one participant reported complete suppression, whereas all others reported perceptual intensities between 1 and 8 (out of max 10 points on the VAS) despite the maximal tolerable noise volume being used.

While it may be feasible to completely suppress concurrent auditory stimulation by applying noise masking at very high sound pressures, we doubt that TMS-related inductive electric stimulation of peripheral sensory and motor axons can be effectively suppressed given the biophysics of TMS. The fast-conducting myelinated peripheral axons passing through the tissue in close proximity to the induced electric filed are readily excitable by TMS (Siebner et al., 1999), and these nerves are exposed to a much larger electric field than the cortex because they are located much closer to the coil. Since myelinated fast-conducting sensory trigeminal fibers are present in parasagittal parts of the dura mater (Lv et al., 2014), concurrent stimulation of dural trigeminal nerve fibers may also contribute significantly to the TEPs. Notably, these nerve fibers are not effectively stimulated by bipolar electric cutaneous stimulation due to the poor electric conductivity of the skull, so that not even our realistic sham condition would be able to control for those responses.

One pioneering TMS-EEG study used electric stimulation of the scalp and did not observe any somatosensory evoked cortical potentials (Paus et al., 2001), yet did neither report the precise stimulation area nor any electric artifact removal procedures. Moreover, it has been argued that SEPs should be located contralateral to stimulation (Du et al., 2017; Paus et al., 2001), concluding that the TEP was unaffected in the absence of a contralateral SEP. However, studies evoking SEPs by face stimulation (including stimulation of the trigeminal nerve) have consistently reported bilateral EEG potentials for both mechanical and electric stimulation of the face (Bennett and Jannetta, 1980; Hashimoto, 1988). While amplitudes were greater contralateral to stimulation, the authors emphasized that the response was bilateral (in contrast to those evoked by afferent nerve stimulation at the wrist, but in agreement with the cortical response distribution of face and head stimulation first reported by Penfield (1937)), and pointed out that the ipsilateral response was heavily contaminated by both muscle and stimulation artefacts. We were able to record ipsilateral SEPs by using a ground electrode near the target site, reverting stimulation polarity after half of the stimulation block to cancel out the electric artefact during averaging, and applying an exponential fitting procedure to subtract the artifact. These procedures revealed an early sham-evoked potential peaking already at a latency of ~25 ms after frontal and parietal sham stimulation. This early potential most likely reflects a cortical SEP component evoked by stimulation of somatosensory trigeminal neurons (Stohr and Petruch, 1979).

### Impact of stimulation intensity

Electric field calculations revealed that the real TMS condition resulted in a highly focal stimulation of the target region which can be attributed to the fact that we used a small figure-of-eight coil with a winding diameter of only 45 mm. Electric field estimations further revealed that focal TMS induced electric fields well above 40-50 V/m in the crown of the targeted gyrus. The induced electric gradient in the cortex is comparable to the values that have been estimated in previous TMS-EEG studies and thought to be well above the threshold for the transcranial induction of cortical responses (Casali et al., 2010; Casali et al., 2013; Massimini et al., 2005; Rosanova et al., 2009; Sarasso et al., 2015). Therefore, we are confident that our real TMS conditions were physiologically effective, inducing a highly focal electric field that was sufficient to evoke action potentials in the targeted cortex. Accordingly, the amplitudes of the TEPs were well within the range of previous TMS-EEG studies on healthy conscious individuals, ranging from 2 to 6 μV (Herring et al., 2015; Kahkonen et al., 2005; Kerwin et al., 2018; Komssi et al., 2004; Noda et al., 2016; Premoli et al., 2014a; Rogasch et al., 2014). In contrast, induced electric field strength achieved by the corresponding sham conditions was well below threshold intensity and thus it is unlikely that it evoked action potentials in the cortical target region.

It is worth pointing out that previous studies have induced stronger electric fields in the cortex with larger figure-of-eight shaped coils, resulting in local responses with higher amplitudes (> 6 μV) (Casarotto et al., 2010; Fecchio et al., 2017; Rosanova et al., 2009). It is conceivable that the relative contribution of the transcranially-evoked response is higher at stronger stimulus intensities and with larger winding of coils (which would stimulate a larger area of the scalp) with the multisensory contribution reaching saturation. However, since concurrent multisensory stimulation will also increase with TMS intensity and with non-focality of the stimulation coil, the stimulus-response relationships for both the transcranially- and peripherally-induced EEG responses need to be systematically characterized in future studies.

## Conclusion

Even though our realistic somatosensory-auditory sham stimulation was not optimally matching the auditory and somatosensory perception of real TMS, we demonstrated substantial similarities between real TMS and sham evoked EEG responses, both at short and long latencies for two cortical target sites. In most experimental settings, it cannot be guaranteed that auditory noise masking and foam padding are sufficient to fully remove any auditory and somatosensory evoked potential in TMS-EEG studies. Therefore, we conclude that the remaining non-transcranial evoked potentials need to be controlled for by multisensory sham conditions. The realistic multisensory sham condition needs to be carefully designed and adjusted to the specific stimulation setup and should match as closely as possible the multisensory stimulation features of real TMS. This should include a systematic psychophysical assessment and comparison of the individually experienced somatosensory and auditory perception of real and sham TMS.

## Acknowledgements

This project has received funding from the following sources: the Capital region of Denmark (Region H, grant number R135-A4841), the Lundbeck Foundation (grant numbers R126-2012-12422, R59-A5399 and R118-A11308), the Danish Council for Independent Research, section of Medical Sciences (DFF-FSS, grant number DFF - 1331-00172) and the Novo Nordisk Foundation (grant number NNF14OC0011413).

